# A developmental ontology for the colonial architecture of salps

**DOI:** 10.1101/2023.09.04.555288

**Authors:** Alejandro Damian-Serrano, Kelly R. Sutherland

**Affiliations:** University of Oregon, Department of Biology, Institute of Ecology and Evolution. 473 Onyx Bridge, 5289 University of Oregon, Eugene, OR 97403-5289

**Keywords:** salps, coloniality, ontology, development, colonial architecture

## Abstract

Colonial animals are composed of clonal individuals that remain physically connected and physiologically integrated. Salps are urochordates with a dual life cycle including an asexual solitary stage that buds sexual colonies composed of jet-propelling zooids that efficiently swim together as a single unit by multi-jet propulsion. Colonies from different species develop distinct architectures characterized by their zooid arrangement patterns, but this diversity has received little attention. Thus, these architectures have never been formally defined using a framework of variables and axes that would allow comparative analyses. We set out to define an ontology of the salp colony architecture morphospace and describe the developmental pathways that build the different architectures. To inform these definitions, we collected and photographed live specimens of adult and developing colonies through offshore SCUBA diving. Since all salp colonies begin their development as a transversal double chain, we characterized each adult colonial architecture as a series of developmental transitions, such as rotations and translations of zooids, relative to their orientation at this early shared stage. We hypothesize that all adult architectures are either final or intermediate stages within three developmental pathways towards either bipinnate, cluster, or helical forms. This framework will enable comparative studies on the biomechanical implications, ecological functions, evolutionary history, and engineering applications of the diversity of salp colony architectures.

## Introduction

Salps (Chordata: Tunicata: Thaliacea: Salpida) are marine pelagic urochordates that filter-feed on phytoplankton and bacteria. The salp life cycle (Fig. 1) consists of a solitary stage (oozooid) that asexually buds colonies of the aggregate stage (blastozooids) along a ventral projection (stolon). Aggregate blastozooids are protogynous and can sexually reproduce, brooding embryonic solitary oozooids in a placenta as females (Bone, 1998). While solitary oozooids move using single-jet propulsion (such as solitary medusae), salp aggregate colonies move in an integrated, coordinated manner through multi-jet propulsion.

**Figure 1.**
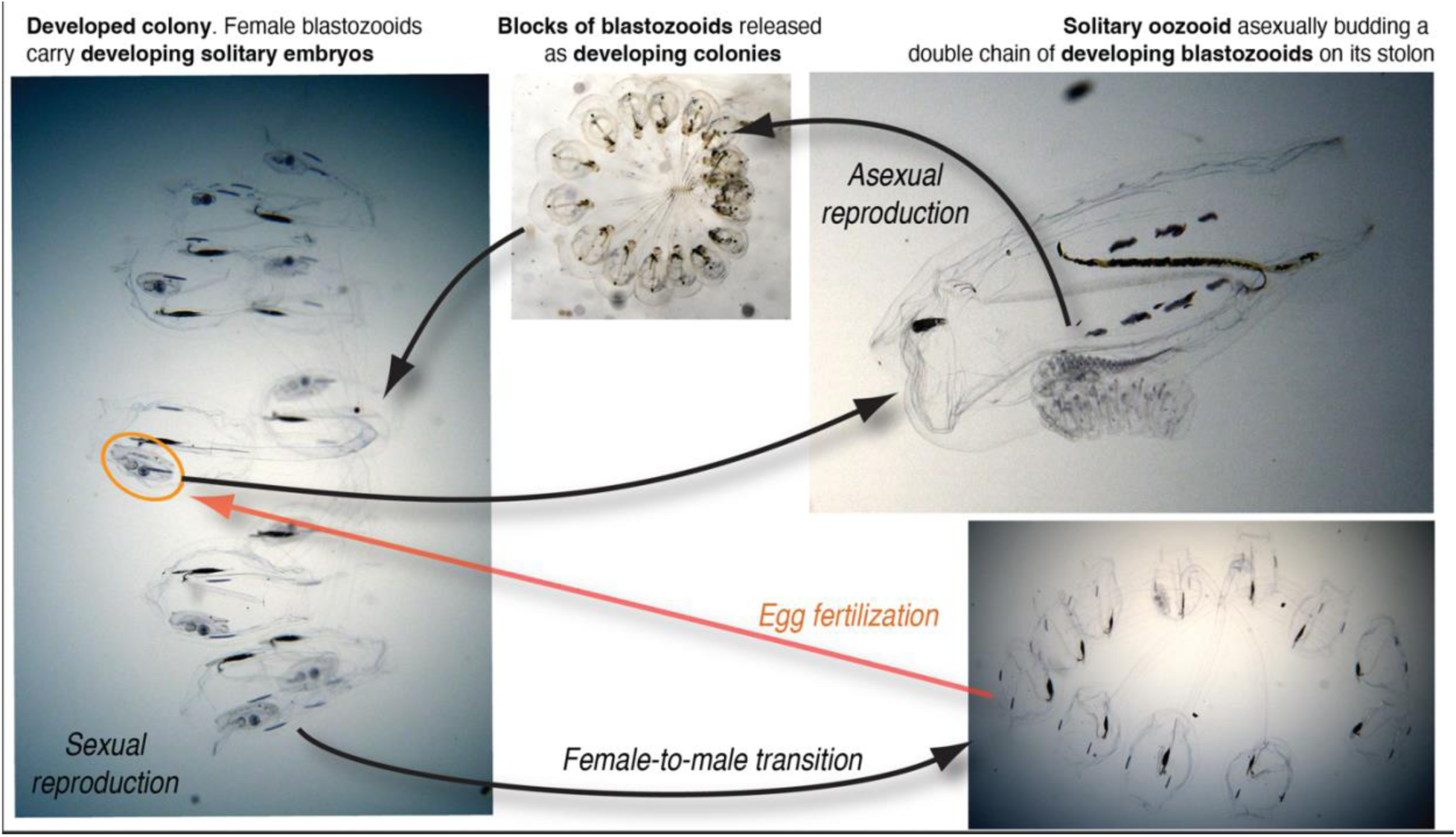
Salp life cycle using the species *Cyclosalpa sewelli* as an example. Frame captures from brightfield in situ videos by Brad Gemmell following method from Colin et al. 2022.

Compared to other multijet colonies, e.g., siphonophores or pyrosomes, salps present a much broader set of architectural configurations among free-swimming colonial animals (Madin, 1990). Salp colony architectures vary across the 48 described species of salps, and include transversal chains, oblique chains, linear chains, whorls, clusters, and helical solenoids (Fig. 2). This diversity and complexity in the arrangements of zooids across species represent a morphological phenotype above the individual’s morphology. A familiar analogy would be the variation in the quaternary structure of proteins as an emergent property of lower-level structural changes. While these colonial architectures look radically different from one another, all species have one early developmental stage in common where the stolon of the solitary progenitor segments into a double chain of paired chiral zooids arranged in a transversal double chain (Bone, 1998).

**Figure 2.**
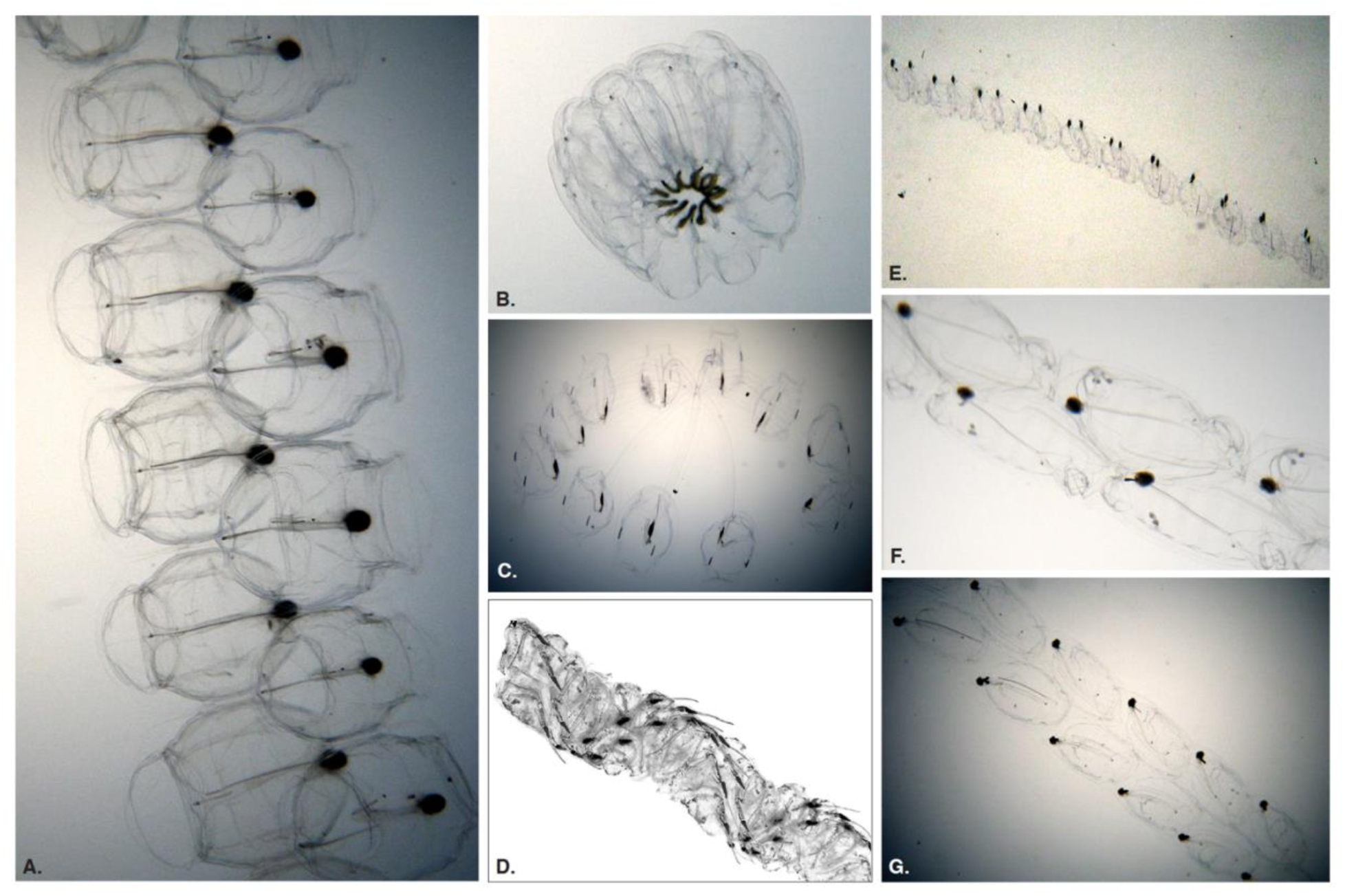
Adult salp colonies representing every distinct colonial architecture observed across all salp species. (A) transversal chain (*Pegea* sp.), (B) whorl (*Cyclosalpa affinis*), (C) cluster (*Cyclosalpa sewelli*), (D) helical chain (*Helicosalpa virgula* (Vogt, 1854), photograph by Nils Aukan), © oblique chain (*Thalia longicauda* (Quoy & Gaimard, 1824)), (F) bipinnate chain (*Ritteriella amboinensis* (Apstein, 1904)). Frame captures A, B, C, E, F, G from brightfield in situ videos by Brad Gemmell.

As the young colony is released from the solitary, the zooids grow rapidly in size, develop their anatomical features, and in most species, they shift their arrangement in the colony into the different architectures we observe in adults. While these colony architectures have been described qualitatively (Madin, 1990), they have received little attention in the past three decades and lack a formal definition. The primary gaps of knowledge include a breakdown of the traits that define the ontology of these architectures, a quantitative framework to measure those traits, and a detailed comparison of the developmental processes that give rise to the different architectures. An ontology is defined as “a set of concepts and categories in a subject area or domain that shows their properties and the relations between them” (Oxford Languages, 2023). In biology, ontologies serve as conceptual frameworks to designate categories, identities, and relationships of parts and variations in complex systems (Bard & Rhee 2004). In this context, we refer to an ontology of salp colony architectures as the categorization of the forms, the description of their characteristics, the definition of their ontogenetic relationships, and their relationships to geometric transformations.

Here we aim to leverage the shared earliest stage in their colonial development to (1) define a set of homologous axes, variables, and planes of observation in all salp colonies, (2) map the different architectures based on (1), and (3) define a hierarchical classification of the distinct types and degrees of developmental translations and rotations of the zooids. The ultimate goal is to enable comparative analyses of variation in zooid arrangements between and within architectures. Using this framework, comparative studies will be able to investigate the biomechanical implications, ecological functions, evolutionary history, and engineering applications of the extant architectonic diversity of salp colonies. Moreover, this work will shed light on the broader design space of clonal coloniality among animals.

## Materials and Methods

We observed and collected live specimens of both adult salp blastozooid colonies and developing colonies in the stolons of solitary salp oozooids. These specimens were collected while SCUBA diving untethered from a small vessel off the coast of Kailua-Kona (Hawai’i Big Island, 19°42’38.7" N 156°06’15.8" W), at an offshore location with a bottom depth of over 2000m. Some dives were conducted during the day, where we encountered most of the specimens of *Iasis cylindrica* (Cuvier, 1804), *Pegea* sp., *Cyclosalpa affinis* (Chamisso, 1819), and *Brooksia rostrata* (Traustedt, 1893). The rest of the species included in this study were collected during night dives when many salps perform diel vertical migration to shallower depths. We collected and photographed blastozooids across 22 salp species. In addition, we supplemented our gaps in taxon sampling using underwater photos and videos of live salps from previous expeditions and from online sources. Salps were identified to the species level using stereo microscopy to examine the dorsal tubercules and muscle band distributions, following the taxonomic keys in Yount (1954), van Soest (1974), Godeaux (1998), as well as in Esnal & Daponte (2009). See SM Table 1 for a list of the specimens observed within each species and architectural type.

After the dive, salp specimens were anesthetized in 0.2% MS222 buffered with sodium bicarbonate in seawater to facilitate photography. Developing stolons were dissected from the anesthetized solitary oozooid before photographing. We photographed anesthetized adult and developing blastozooid colonies in glass crystallization dishes with a black background using a Canon 6D DSLR camera with a 35mm lens mounted on an inverted tripod used as a copy stand. Specimens were photographed from different orientations relative to the constituent zooids’ bilateral symmetry (oral, aboral, dorsoventral, and lateral), with a ruler in the frame for scale reference.

From these images, we examined the colonial arrangement from the earliest stage of stolon development to adulthood. In some taxa, the temporal axis of blastozooid development can be observed spatially in a continuous gradation of blastozooid development (Fig. 3). This is the case in *Cyclosalpa* spp. (Fig. 3E-F), *Brooksia* spp., *Soestia* spp., and *Helicosalpa* spp. (Fig. 3G). In other taxa, the temporal axis of blastozooid development can also be observed spatially in discretely segmented cohort blocks with synchronous development within each block, such as in *Salpa* spp., *Ritteriella* spp. (Fig. 3D), and *Thalia* spp. Other taxa, however, produce only a single cohort block with synchronous development, such as in the case of *I. cylindrica* (Fig. 3C)*, Thetys vagina* Tilesius, 1802 (Fig. 3B)*, Pegea* spp. (Fig. 3A), and *Traustedtia multitentaculata* (Quoy & Gaimard, 1834). We examined the development of the blastozooid chain in these taxa by keeping the solitaries alive in seawater and observing the developmental transitions overnight.

**Figure 3.**
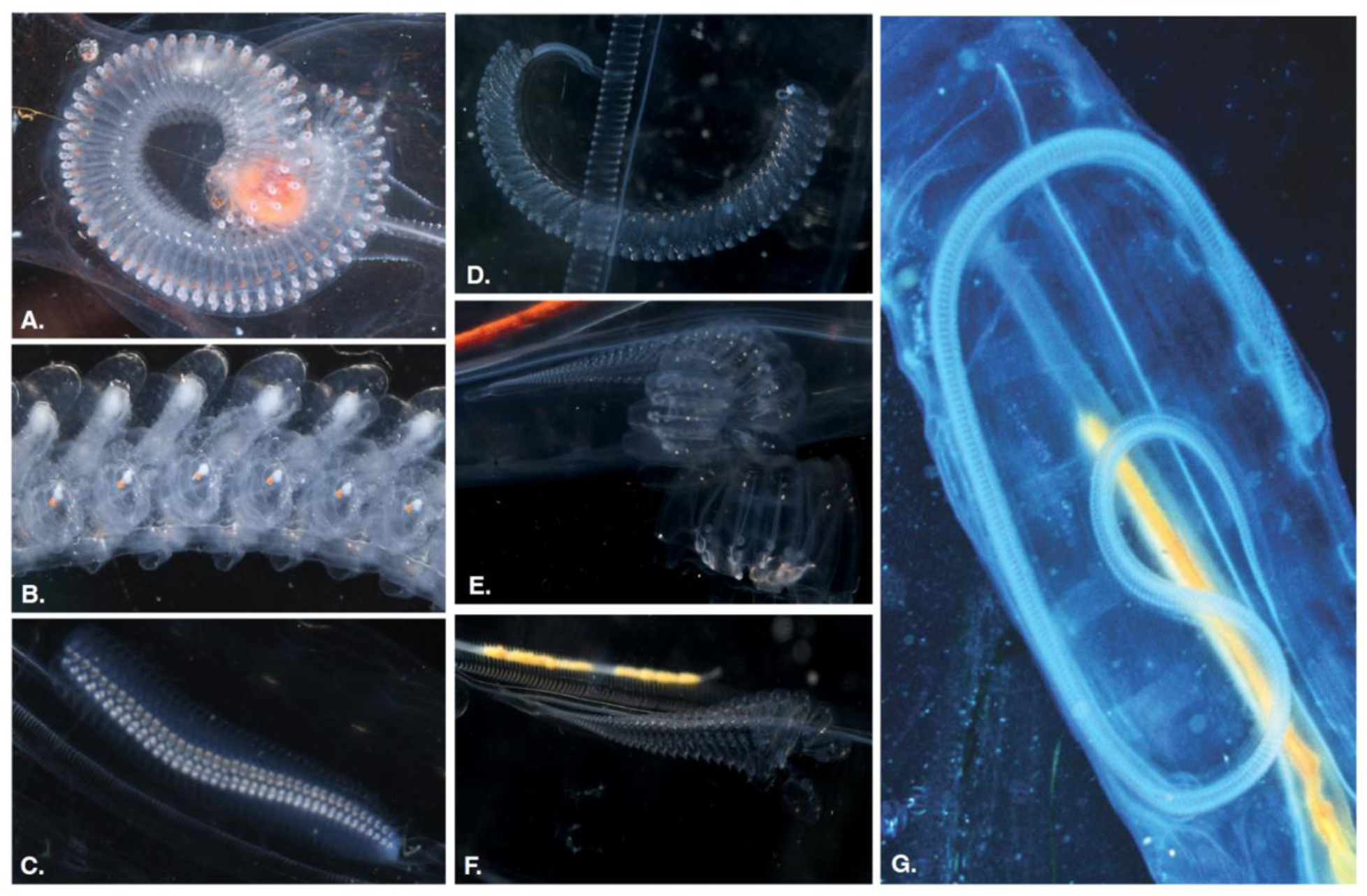
Developing blastozooid colonies produced by the budding stolons of solitary oozooids across different salp architectures. (A) transversal chain buds (*Pegea* sp.), (B) oblique chain buds (*Thetys vagina*), (C) linear chain buds (*Iasis cylindrica*), (D) bipinnate chain buds (*R. amboinensis*), (E) whorl buds (*C. affinis*), (F) cluster buds (*Cyclosalpa polae*), (G) helical chain buds (*H. virgula*, photograph by David Wrobel).

## Results

### Defining the observation framework

The arrangement and relative orientation of blastozooids in different colony architectures present a 3-dimensional problem, where the axes and angles of reference shift in ways that are challenging to compare from a single viewpoint. Using the transversal double-chain architecture found in the earliest developmental stage of every species (as well as in adult colonies of *Pegea* spp. and *Traustedtia* spp.), in addition to the bilateral symmetry of salp blastozooids, we defined three orthogonal axes and their corresponding normal planes (Fig. 4). These are: (1) The dorsoventral axis of the colony is defined as the axis parallel to the dorsoventral axis of the zooids in the transversal double chain, with a normal (perpendicular) plane of observation corresponding to viewing the dorsal side of the zooids on either side of the transversal double chain. (2) The oral-aboral axis is defined as the axis parallel to the oral-aboral axis of the zooids in the transversal double chain, with a perpendicular plane of observation corresponding to viewing the oral or aboral end of the zooids on either the frontal or rear side of the transversal double chain. The zooid oral-aboral axis of each zooid is defined as the line parallel to the endostyle. (3) The stolon axis is defined as the axis of chain growth parallel to the stolon, perpendicular to a plane of observation that corresponds with looking directly at either end of the transversal double chain, with a lateral view of the zooids.

**Figure 4.**
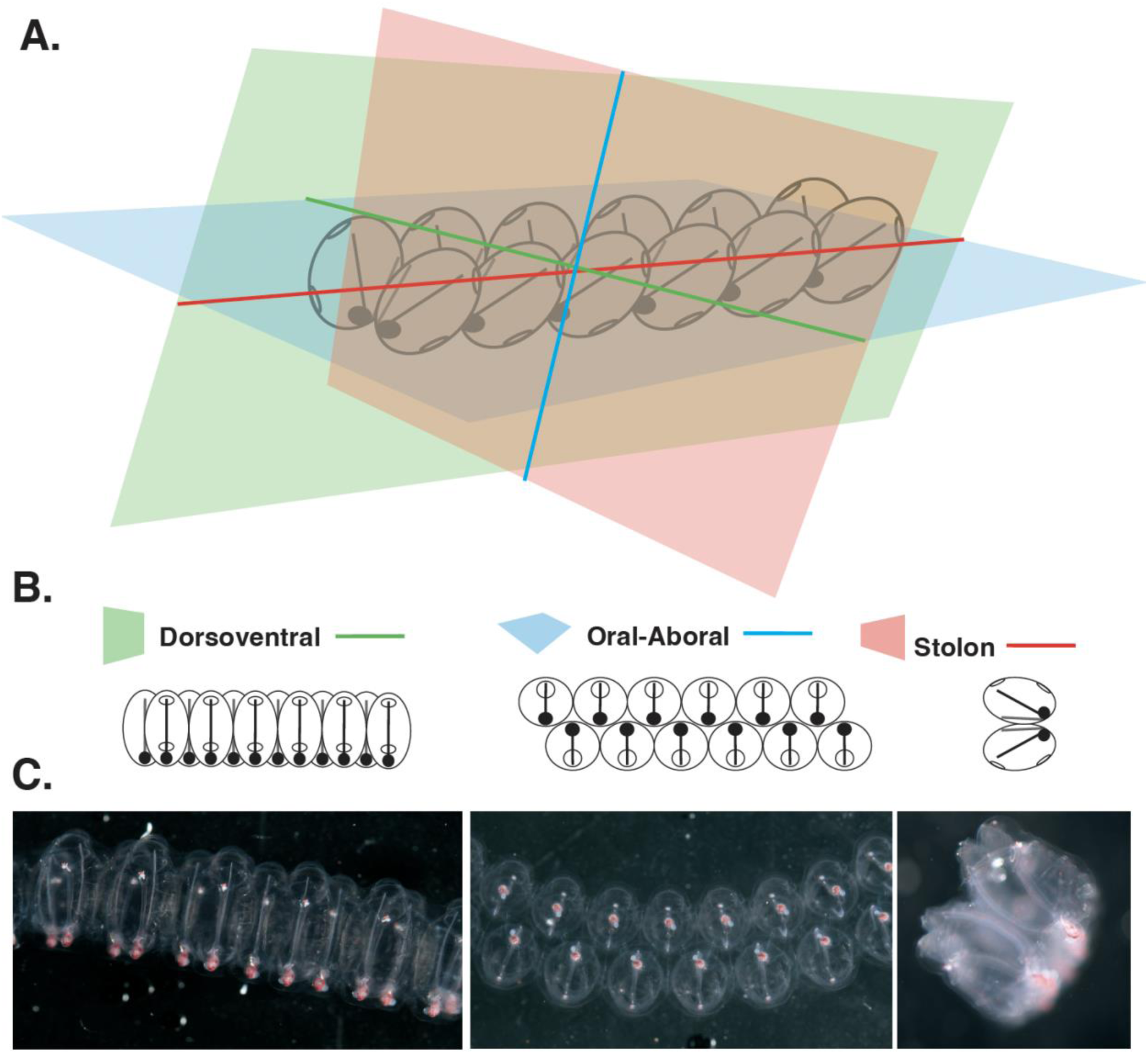
(A) Definition of the homologous universal axes and planes of observation relative to the orientation of the colony in the early transversal chain stage. (B) Diagrams representing the cross-sectional views of a transversal chain from each of these three planes of observation. (C) Photographs of a transversal chain of early-developing *Pegea* sp. blastozooids taken from each of the abovementioned orientations.

With these axes and planes delineated, we can then describe developmental changes in zooid orientation characters relative to these three planes of observation. Based on this universal observation framework, we then defined the following characters: (A) the dorsoventral zooid-stolon angle is the angle formed between the oral-aboral axis of the zooid and the elongation axis on the colony as viewed from the dorsal side of the developing zooids, driving the formation of oblique, linear, and bipinnate chains; (B) the lateral chiral angle, defined as the angle formed between the oral-aboral axes of a pair of chiral zooids as viewed from the zooids’ lateral orientation, driving the formation of bipinnate chains; (C) zooid autorotation, defined as rolling of the zooid around its own oral-aboral axis also driving the formation of bipinnate chains; (D) serial stolon-normal angle, defined as the angle formed between a zooid’s oral-aboral axis and the oral-aboral axis of its lateral neighbor as viewed from one end of the colony, driving the formation of a solenoid double helix chains; (E) peduncle length ratio (the peduncle is an extension of the tunic that connects the zooids to their chiral pair or to the stolon during development, present in most *Cyclosalpa* species), defined as the ratio between the total oral-aboral length of the zooids relative to the longest axis of their peduncle, driving the formation of whorls and clusters; and finally (F) neighbor attachment, defined as direct contact between lateral neighbors, its loss drives the formation of clusters with loosely-attached zooids.

We examined changes in these variables across the development of colonies in different salp species and characterized a developmental ontology of salp colony architecture by first describing the set of developmental transformations that give rise to each architecture, then identifying which intermediate stages in the formation of more derived architectures in some species are equivalent to the adult finalized architectures in other species, to build a process-based hierarchical ontology of the architectures within colonial developmental pathways.

### A developmental ontology of architectural transition pathways

In congruence with Madin (1990), we observed that the earliest stages in the development of salp colonies across all species display a transversal double chain architecture which undergoes subsequent changes towards the adult colonial architecture.

In some taxa, such as *Pegea* spp. (with adult and developing forms examined in this study) and *Traustedtia* spp. (observed in photographs from other divers), we noticed that the transversal double-chain architecture remains unchanged throughout the growth and development of the blastozooids in the chain. This architecture is characterized by a dorsoventral zooid-stolon angle of ∼90°, with ventral attachment to the chiral neighbor and lateral attachment to the lateral neighbors (Fig. 2A). In the field, we observed these chains moving parallel to the oral-aboral axis of their zooids, at an angle orthogonal to the length of the chain. Often, we find colonies of *Pegea* species moving in a coiled formation, where the transversal chain is curled up on the oral-aboral-normal plane. Other species we examined do not retain this developmentally basal architecture, but instead modify the orientation, rotation, and position of the zooids relative to each other and the axis of the chain during development (Fig. 5). Our observations on the developmental series of salp species revealed intermediate stages between the initial transversal arrangement and the final adult arrangement (terminal architecture). In the case of the most elaborate terminal architectures (such as bipinnate chains, linear chains, and clusters) their intermediate stages shared the diagnostic characteristics of other simpler terminal architectures. We summarized the developmental relationships between architectures as three hypothesized developmental transition pathways (depicted in Figure 6), each characterized by specific zooid rearrangement processes.

**Figure 5.**
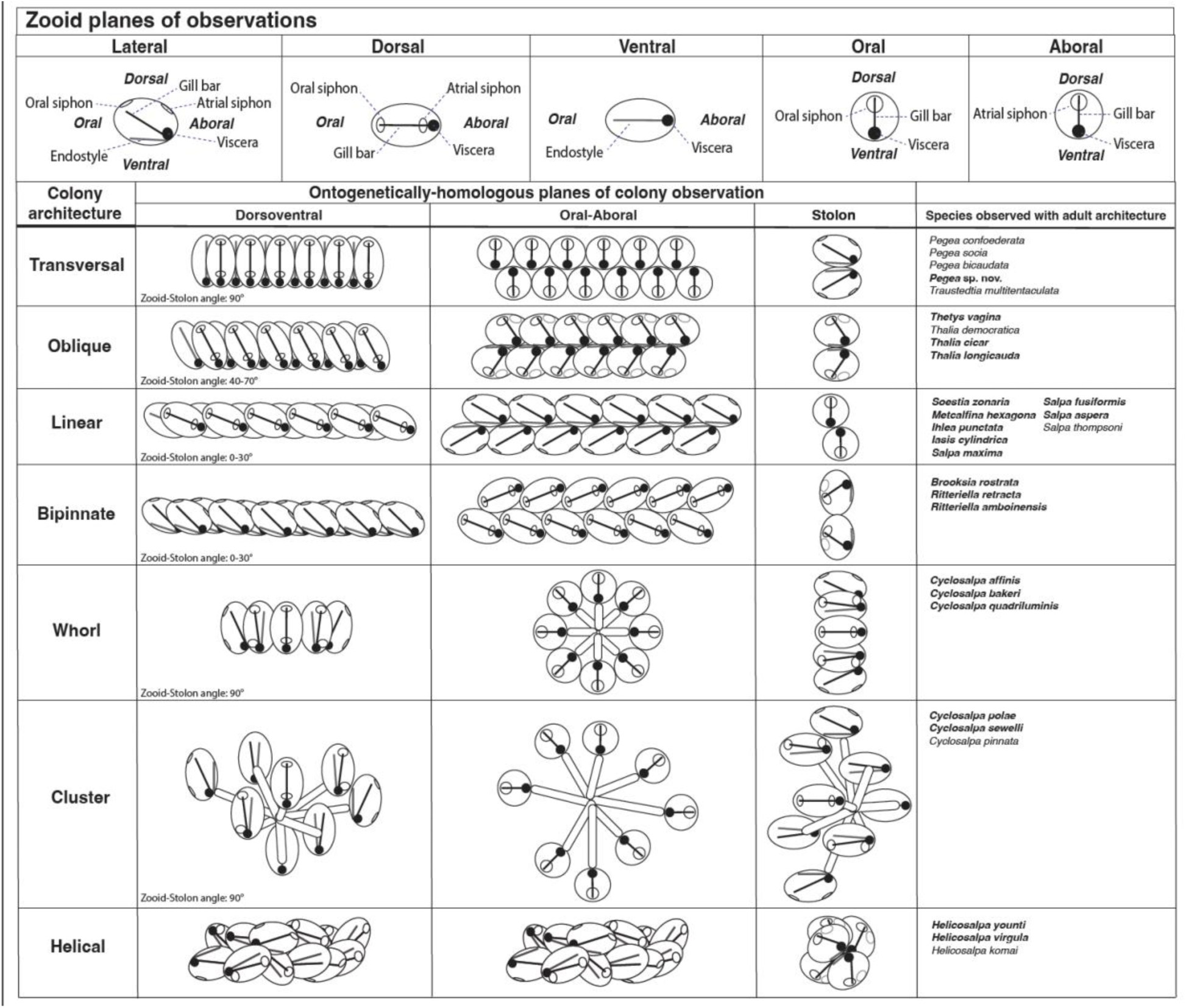
Sketches of individual zooids and adult colonies representative of every architecture as viewed from each plane of observation, with key structures labeled. The rightmost column lists the species in which these terminal architectures were observed.

**Figure 6.**
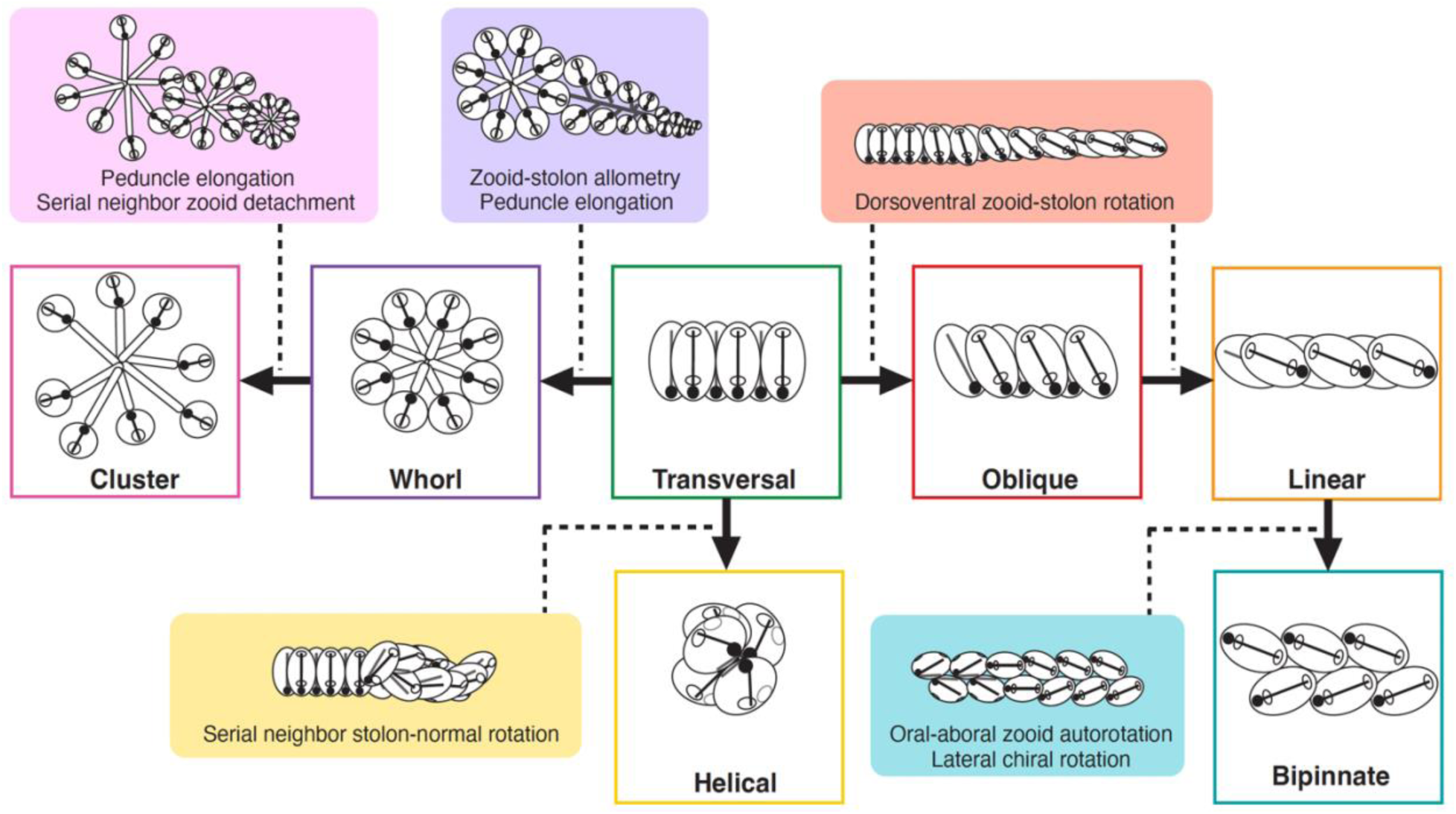
Hypothesized developmental transitions (solid arrows) and mechanisms involved (solid color boxes associated with each transition by a dotted line) leading to the different adult blastozooid colony architectures (white boxes with colored outlines). All developmental mechanisms are based on observations of colonial development, except those involved in the bipinnate architecture, which are hypothetical and based on comparisons between adult architectures. The transversal architecture is found in the earliest developmental stage of every species as well as in the adult stage of some species and has been hypothesized to be ancestral to all salps (Madin 1990).

First there is the hypothesized pathway that leads to the architectures found in *Cyclosalpa* spp. These blastozooid chains continue their development in a transversal arrangement (dorsoventral zooid-stolon angle of 90°) but grow peduncles that separate the zooids from the stolon attachment point and from their chiral ventral neighbor. Moreover, discrete sets of budding salps bundle together, where the attachment points of their peduncles remain attached to a central point and detached from other sets. These radial aggregations (whorls) are formed by two chiral, bilaterally symmetrical, semicircular sets of zooids (Ritter & Johnson 1911). In the first stage of the transformation, we observe the formation of the whorl architecture (e.g., Fig. 3E). The zooids in this architecture are packed together tightly in a wheel shape due to the short peduncles (Fig. 2B). We found this terminal architecture and observed its development in some *Cyclosalpa* species such as *C. affinis, C. quadriluminis* Berner, 1955, and *C. bakeri* Ritter, 1905. In *C. affinis*, we saw these whorls remaining attached to each other for a longer time than in other species and adult whorls can often be found conjoined. We hypothesize that a further (more developmentally derived) stage in this transformation is the cluster architecture, where after reaching the ‘whorl’ intermediate stage, the peduncles grow longer to the point that zooids are no longer attached laterally to each other (Fig. 2C) and can freely bob around and end up arranged in hemispherical or spherical sets. We found the cluster terminal architecture in other *Cyclosalpa* species such as *C. sewelli* Metcalf, 1927*, C. pinnata* (Forskål, 1775) (observed in the field but not photographed for this study), and *C. polae* Sigl, 1912. We observed the development of cluster architectures from a transversal stage in *C. polae* and *C. sewelli*. These cluster bundles appeared to contain many more zooids than those in whorls. From our observations of the development of *C. sewelli*, we were not able to discern whether they ever go through a distinct ‘whorl’ stage the way that *C. polae* definitively does. The developing cluster and whorl colonies we observed all went through a similar zooid-stolon allometry, though we were unable to determine whether the serial neighbor zooid detachment (Fig. 6) occurs before or after the release of the radial aggregations.

Second, there is the hypothesized pathway leading to the helical double-chain architecture in *Helicosalpa* spp. (Fig. 2D). While we did collect and examine adult colonies of *H. virgula* and *H. younti*, we did not encounter any *Helicosalpa* oozooids with developing colonies in which to observe the developmental transition. However, we could observe their development in a photograph by David Wrobel (Fig. 3G), and in Figure 2 of Ringvold et al. (2020). Like the other salps we observed, these double chains are budded in a transversal arrangement, but subsequently undergo stolon torsion into a solenoid shape (Fig. 3G), where zooids become angled relative to their chiral pair to accommodate this conformation. Then there is the hypothesized pathway leading to streamlined linear chains through the alignment of zooid orientations to the axis of the stolon during development. We observed the development of linear chains from transversal stages in *Iasis cylindrica* and noticed they start with a partial dorsoventral rotation of zooids at an angle within the range of oblique terminal architectures (Fig. 3B), which is the final form for species like *Thetys vagina* (as mentioned in Madin 1990) and *Thalia* spp. (at least in *T.longicauda* and *T. cicar* examined in this study, as well as in *T. democratica* observed in other sites by the authors, see example in Fig. 2E), with zooid-stolon angles between 50-60°. Therefore, we hypothesize that the swimming direction of these oblique colonies is closely aligned (but not perfectly parallel) with the stolon axis. In subsequent stages of the development of linear colonies, we observed this dorsoventral torsion going further toward near-complete alignment (15-30°) of the oral-aboral axis of the zooids to the axis of the stolon. We observed a terminal linear architecture in *Ihlea punctata, Iasis cylindrica, Metcalfina hexagona*, *Soestia zonaria* (Pallas, 1774), and several *Salpa* species. In the species *S. zonaria*, we find the most extreme version of this architecture, with zooid-stolon angles close to 0° (Fig. 2F). Finally, based on similarities between the zooid orientations between bipinnate and linear terminal architectures, we hypothesize that the bipinnate architecture (which we observed in *Brooksia rostrata, Ritteriella amboinensis,* and *Ritteriella retracta*) represents a derived transformation stage in the linear pathway. We observed that both linear and bipinnate architectures share the linear dorsoventral alignment of zooids to the stolon. However, adult colonies in the species we observed having the bipinnate architecture additionally displayed a symmetrical outward lateral flare of the aboral ends of zooids in the oral-aboral-normal plane and an autorotation of zooids where the ventral (and dorsal) sides of every zooid are all facing the same side of the colony (Fig. 2G). The developmental transitions in bipinnate species was challenging to observe empirically because *Ritteriella* does not undergo transformation past the oblique stage in colonies retained by the oozooid (Fig. 3D), and in *Brooksia* the transformation occurs at a very small scale in the most proximal and underdeveloped end of the budding colony. It is possible that the order of developmental transitions that lead to the bipinnate morphology differs from the one hypothesized in Figure 6. However, our observations on developing *Ritteriella* colonies indicate that these transformations occur during or after the process of dorsoventral zooid stolon rotation that produces the oblique intermediate form.

Each developmental transition is characterized by variation across specific continuous morphological traits (Fig. 6), which we describe based on the differences in zooid orientation and shape between the initial budding transversal chain stage and the final adult stage supplemented by some observations of the developmental changes. We hypothesize that the transversal-to-whorl transformation is mediated by an increase in the peduncle-to-zooid length ratio and a continuous allometric shift in zooid-to-stolon size as the zooids grow and develop asynchronously along the stolon length. The subsequent whorl-to-cluster transformation also seems to rely on further peduncle elongation but is marked by a loss of neighbor zooid attachment that allows neighboring zooids to bob around freely. The transversal-to-helical pathway (based on observations made on the photograph in Fig. 3G) is characterized by a continuous shift in the serial neighbor stolon-normal angle, where the orientation of neighboring zooids breaks parallelism and starts to offset by a few degrees like stairs in a spiral staircase. The transversal-to-oblique-to-linear pathway appears to be driven solely by changes in the dorsoventral zooid-stolon angle based on our observations. Finally, we hypothesize that the linear-to-bipinnate transformation is driven by an increased oral-aboral chiral angle and zooid autorotation, where the oral-lateral facets of chiral zooids face each other, the aboral ends turn outwards, and their ventral sides face the same side of the colony. However, we were not able to observe the development of bipinnate colonies past an oblique stage, so the order in which these modifications occurred remains unknown.

## Discussion

Ontologies in biology are helpful conceptual tools to characterize, categorize, and compare variation between and within species. We leveraged similarities in the development of salp colonies across species to categorize and geometrically compare the different architectures. From this developmental perspective, we were able to propose a hypothesis-generating ontology for salp colony architecture describing the developmental transitions in the zooid arrangements that lead to the different terminal architectures. Through this ontology, we hypothesize that some terminal architectures are homologous to intermediate stages in the development of other terminal architectures. These ontological definitions and reference frameworks are essential to measure and compare standing variation in colony architecture and its emergent properties between salp species.

One of the most immediate emergent properties of salp colony architecture is the potential implications for locomotion. Different salp colony architectures present different relative orientations of the individual jets to each other and to the overall colony motion axis. In addition, we hypothesize that different architectures differ in how the number of zooids in the colony scales with cross-sectional area relative to motion. These hydrodynamic properties can have further consequences on the locomotory efficiency of different architectures. Swimming in linear salp chains is hypothesized to be more economical due to the reduction of drag (Bone & Trueman 1983). A salp colony is equipped with multiple propelling jets rather than one, which increases its propulsive power. Drag experienced during swimming depends on the total area exposed to the fluid as well as the frontal (motion-orthogonal) projected area (Alexander 1968). Skin drag will increase with the number of zooids in the colony in a predictable manner that is independent of their zooid arrangement. However, frontal drag is drastically reduced in linear chains compared to the sum of each separate zooid (Mackie 1986). We hypothesize that frontal drag will vary across architectures and therefore impact the relative speed attained by each species. In addition to changing the way the frontal area scales with the number of zooids, we hypothesize that architecture may also impact the angles of the jets relative to the axis of colony motion. In siphonophores, the velum of the nectophore is used to orient the jet to prioritize torque or thrust (Sutherland et al 2019). In salps, these orientations are usually fixed in a colony (Sutherland & Weihs 2017), but the angle of the exhalant jets relative to the swimming of the colony will dictate the thrust-to-torque ratio, which will determine their propulsive efficiency.

Understanding the hydrodynamic advantages and implications of each colonial architecture can be valuable beyond basic science since it may yield interesting applications to bioinspired underwater vehicles. Pulsatile jet propulsion is increasingly inspiring underwater vehicle engineering (Mohensi 2006, Yue et al. 2015). Multijet systems comprised of collaboratively interactive propeller units could revolutionize the field of underwater vehicles (Chao et al. 2017, Costello et al. 2015) with designs inspired by gelatinous invertebrates such as salps (Marut 2014, Krummel 2019, Bi et al 2022). Some of these bio-inspired solutions are stimulating novel solutions in the field of soft robotics (Renda et al. 2015, Krummel 2019), as deformable body shapes can augment propulsive forces (Giorgio-Serchi & Weymouth 2017). Understanding the biomechanical underpinnings of the diversity of salp colony architectures would reveal nature’s broadest design space for underwater multi-jet-propelled soft locomotors and their inherent trade-offs.

Another potential contribution of this architectural ontology is the characterization of colonial morphology from a comparative, evolutionary perspective. Salp colony architectures are distributed across the phylogenetic diversity of salp species, but their evolutionary history remains unknown. The two main obstacles to the reconstruction of the evolutionary history of salp colony architecture have been (1) the lack of a framework to compare and characterize variation between architectures, and (2) a phylogenetic tree that resolves the position of every architecture in every lineage where it has evolved. Metcalf (1918) hypothesized phylogenetic relationships among salps based on gut morphology, with *Cyclosalpa* as the most distant relative to other salps due to its linear gut shape. Half a century later, Madin (1974) hypothesized that colonial architecture is phylogenetically conserved and that fast-swimming linear and bipinnate chains are monophyletic. Years later, Tsakogeorga et al. (2009) reconstructed the first molecular phylogeny using 18S sequences to resolve relationships between thaliacean groups, supporting the monophyly of salps. Following this work, Govindarajan et al. (2011) included a more extensive taxon sampling within salps, revealing that salps with linear chain architectures are not monophyletic, and that the transversal architecture in *Pegea* is likely derived, despite having been hypothesized as ancestral (Madin 1990). While this phylogeny included many of the known salp species, it cannot fully resolve the evolutionary history of salp colony architecture since the position of *Pegea* and *Thalia* are poorly resolved and the position of *Helicosalpa* is unknown. A phylogenetic comparative approach to the diversity of colonial architectures will facilitate further research on its evolutionary, ecological, and biomechanical underpinnings. If evolutionary shifts in the architecture of salp chains bring on changes in their locomotory efficiency, it is possible that these shifts are related to different selective pressures such as predatory pressure, habitat nutritional patchiness, or vertical migration behavior.

The research directions outlined above would advance our understanding of salp biology across their species diversity. Salps have attracted significant scientific interest in the past decade since they are essential consumers in oceanic ecosystems that feed on microbial plankton production and can grow explosively following phytoplankton blooms (Henschke et al. 2016). Salp fecal pellets play an important role in the biological carbon pump and are responsible for a large fraction of the biological carbon pump (Decima et al. 2023), responsible for the trapping of gigatons of carbon fixed from the atmosphere into the deep sea and are therefore a key mediator for atmospheric CO_2_ concentration and global change (Buesseler et al. 2020). Many of the salp species that contribute most to this process are vertical migrators that respire and deposit (by defecation and predation) carbon during the day in the mesopelagic zone, after feeding during the night near the surface (Steinberg et al 2023). Many of the vertically migrating salp species (such as *Salpa* spp.) present a linear architecture (Madin et al. 1996) and their migratory behavior varies with colonial development (Henschke et al. 2021). While some of these linear, vertically migrating species have been extensively investigated, the ecology and natural history of the broader diversity of salps remains understudied. Characterizing the relationship between colonial architecture, locomotion, and migratory behavior is key to understanding the ecological implications of shifting salp species compositions and distributions with global change (Lavaniegos & Ohman 2003).

Finally, we believe the colonial ontology proposed here expands our understanding of the development and evolution of colonial animals in general. Colonial animals (modular colonies, not including eusocial colonies) are composed of clonal individuals produced by asexual reproduction that remain physically connected and physiologically integrated (Harvell, 1991). Most animal colonies are arranged with their zooids in parallel to each other forming 2D sheets with one pole, typically the oral end, exposed to the external environment. Some of these topologically simple planar colonies can form complex 3D shapes by folding this sheet. In benthic species, the sheet is often folded around an endogenous skeleton or an object in the environment (e.g., corals, ascidians, millepores). Bryozoans also tend to develop into sheets though also into branching structures. In pelagic species, colonies are free-living and capable of swimming around by the combined (and often coordinated) action of their zooids (Du Clos et al. 2022). Therefore, their shapes are often directional, with a front and a rear end defined by their colonial locomotion (Mackie, 1986). On one hand, pyrosomes (Chordata: Tunicata) use the same 2D-sheet template as their benthic relatives, yet in their case, the sheet grows folded forming a closed-ended tube where all the exhalant flow from the inner side aboral ends of the zooids are canalized to a single jet stream. On the other hand, siphonophores and doliolids typically form 1D colonies with sub-specialized zooid types, where only one or few frontal locomotory zooids (nectophores and nurse zooid, respectively) propel a linear colony with non-swimming zooids dragging behind. Siphonophore (Cnidaria: Hydrozoa) colonies can be topologically complex in benthic rhodaliids or in the pleustonic Portuguese Man-o-war, but most planktonic free-swimming siphonophore colonies have their zooids arranged bi-serially or mono-serially along a stem (Mackie et al., 1988). Among them, physonect siphonophores bear multiple nectophores (swimming bodies) that propel the colony through multijet propulsion (Sutherland et al. 2019) in a similar fashion to linear salp colonies. Compared to siphonophores or pyrosomes, salps present a much broader set of architectural configurations among free-swimming colonial animals (Madin, 1990), thus expanding the boundaries of our known design space for both form and function of coloniality in the pelagic realm.

## Acknowledgments

We would like to thank the crew of Aquatic Life Divers and Kona Honu Divers for their help and support in hosting our offshore diving operations. We would also like to thank Marc Hughes, Kevin Du Clos, Jeff Milisen, Brad Gemmell, Sean Colin, Jack Costello, Rebecca Gordon, Matt Connelly, Clint Collins, Paul Richardson, and Anne Thompson for their assistance during diving and photography operations in the field. This research was funded by the Gordon and Betty Moore Foundation (grant 8835) and the Office of Naval Research (N00014-23-1-2171).

## Ethical Care Considerations

Our specimen collection and protocol were compliant with all local regulations. Since no vertebrates or cephalopods were involved, we did not need oversight from an animal care board.

## Data Accessibility

All photographs will be made available in a Dryad repository.

## Supplementary Data

**SM Table 1.**
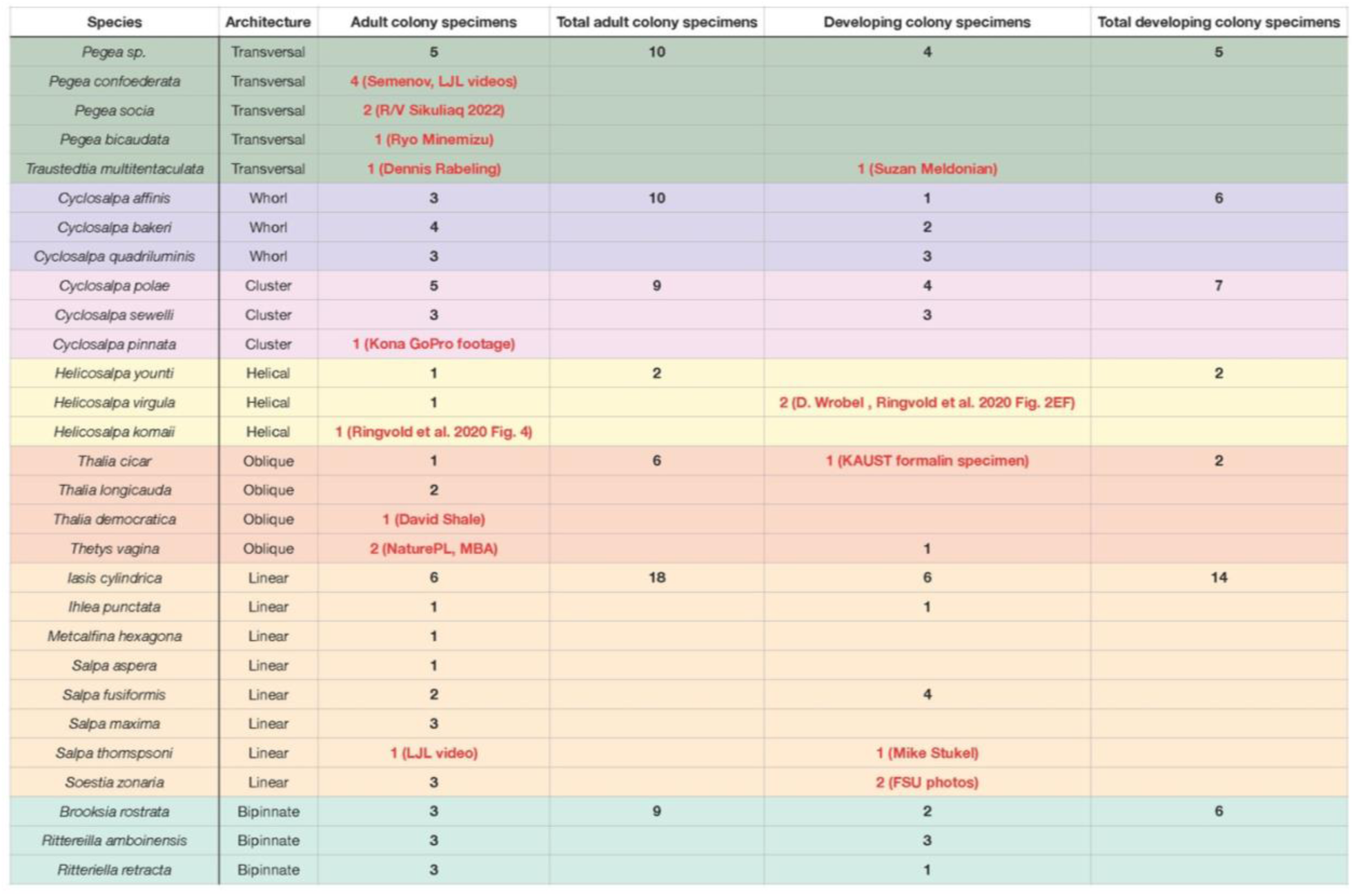
Number of images examined from salp specimens of each life cycle stage and species. Specimens collected visa SCUBA diving off Kona are shown as numbers in black. Other images from different sources are highlighted in red.

## Literature Cited

Alexander, W., 1968, September. A discussion of governing decelerator performance and design parameters in the supersonic flight regime. In 2^nd^ Aerodynamic Deceleration Systems Conference (p. 938).

Bard, J. B., & Rhee, S. Y., 2004. Ontologies in biology: design, applications and future challenges. Nature reviews genetics, 5(3), 213–222.

Bi, X., Tang, H., & Zhu, Q., 2022. Feasibility of hydrodynamically activated valves for salp-like propulsion. Physics of Fluids, 34(10), 101903.

Bone, Q., 1998. The biology of pelagic tunicates.

Bone, Q. and Trueman, E.R., 1983. Jet propulsion in salps (Tunicata: Thaliacea). Journal of Zoology, 201(4), pp.481–506.

Buesseler, K. O., Boyd, P. W., Black, E. E., & Siegel, D. A., 2020. Metrics that matter for assessing the ocean biological carbon pump. Proceedings of the National Academy of Sciences, 117(18), 9679–9687.

Chao, S., Guan, G., & Hong, G. S., 2017, September. Design of a finless torpedo shaped micro AUV with high maneuverability. In OCEANS 2017-Anchorage (pp. 1-6). IEEE.

Colin, S. P., Gemmell, B. J., Costello, J. H., & Sutherland, K. R. (2022). In situ high-speed brightfield imaging for studies of aquatic organisms. Protocolsio.

Costello, J. H., Colin, S. P., Gemmell, B. J., Dabiri, J. O., & Sutherland, K. R., 2015. Multi-jet propulsion organized by clonal development in a colonial siphonophore. Nature communications, 6(1), 8158.

Décima, M., Stukel, M. R., Nodder, S. D., Gutiérrez-Rodríguez, A., Selph, K. E., Dos Santos, A. L., … & Pinkerton, M., 2023. Salp blooms drive strong increases in passive carbon export in the Southern Ocean. Nature communications, 14(1), 425.

Du Clos, K. T., Gemmell, B. J., Colin, S. P., Costello, J. H., Dabiri, J. O., & Sutherland, K. R., 2022. Distributed propulsion enables fast and efficient swimming modes in physonect siphonophores. Proceedings of the National Academy of Sciences, 119(49), e2202494119.

Esnal, G. B. & M. C. Daponte, 1999. Salpida. In Boltovskoy, D. (ed.), South Atlantic Zooplankton. Backhuys Publishers, Leiden: 1423–1444.

Giorgio-Serchi, F., & Weymouth, G. D., 2017. Underwater soft robotics, the benefit of body-shape variations in aquatic propulsion. In Soft Robotics: Trends, Applications and Challenges: Proceedings of the Soft Robotics Week, April 25-30, 2016, Livorno, Italy (pp. 37-46). Springer International Publishing.

Godeaux, J., 1998. The relationships and systematics of the Thaliacea, with keys for identification. In: Q. Bone (ed.), The Biology of Pelagic Tunicates: 273–294. Oxford Univ. Press. Oxford.

Govindarajan, A.F., Bucklin, A. and Madin, L.P., 2011. A molecular phylogeny of the Thaliacea. Journal of Plankton Research, 33(6), pp.843–853.

Harvell, C.D., 1991. Coloniality and inducible polymorphism. The American Naturalist, 138(1), pp.1–14.

Henschke, N., Everett, J. D., Richardson, A. J., & Suthers, I. M., 2016. Rethinking the role of salps in the ocean. Trends in Ecology & Evolution, 31(9), 720–733.

Henschke, N., Cherel, Y., Cotté, C., Espinasse, B., Hunt, B. P., & Pakhomov, E. A., 2021. Size and stage specific patterns in *Salpa thompsoni* vertical migration. Journal of Marine Systems, 222, 103587.

Krummel, G. M., 2019. Locomotion and Control of Cnidarian-Inspired Robots (Doctoral dissertation, Virginia Tech).

Lavaniegos, B. E., & Ohman, M. D., 2003. Long-term changes in pelagic tunicates of the California Current. Deep Sea Research Part II: Topical Studies in Oceanography, 50(14-16), 2473–2498.

Mackie, G.O., 1986. From aggregates to integrates: physiological aspects of modularity in colonial animals. Philosophical Transactions of the Royal Society of London. B, Biological Sciences, 313(1159), pp.175–196.

Mackie, G.O., Pugh, P.R. and Purcell, J.E., 1988. Siphonophore biology. In Advances in Marine biology (Vol. 24, pp. 97-262). Academic Press.

Madin, L. P., 1974. Field studies on the biology of salps, University of California, Davis (pg. 1 208) PhD Thesis.

Madin, L.P., 1990. Aspects of jet propulsion in salps. Canadian Journal of Zoology, 68(4), pp.765–777.

Madin, L. P., Kremer, P., & Hacker, S., 1996. Distribution and vertical migration of salps (Tunicata, Thaliacea) near Bermuda. Journal of Plankton Research, 18(5), 747–755.

Metcalf, M. M. and Bell, M. M., 1918. The Salpidae: a taxonomic study (Vol. 2). US Government Printing Office.

Sutherland, K. R., Gemmell, B. J., Colin, S. P., & Costello, J. H., 2019. Propulsive design principles in a multi-jet siphonophore. Journal of Experimental Biology, 222(6), jeb198242.

Marut, K. J., 2014. Underwater Robotic Propulsors Inspired by Jetting Jellyfish (Doctoral dissertation, Virginia Tech).

Mohensi, K., 2006. Pulsatile vortex generators for low-speed maneuvering of small underwater vehicles. Ocean Eng. 33, 2209–2223.

Oxford Languages, 2023. Oxford Languages and Google – English. Oup. Com. https://languages.oup.com/google-dictionary-en/

Renda, F., Serchi, F. G., Boyer, F., & Laschi, C., 2015. Structural dynamics of a pulsed-jet propulsion system for underwater soft robots. International Journal of Advanced Robotic Systems, 12(6), 68.

Ringvold, H., Hatlevik, A., Hevrøy, J., Hughes, M. and Aukan, N., 2020. Encounters with the rare genus Helicosalpa (Chordata, Thaliacea, Salpida), using citizen science data. Marine Biology Research, 16(5), pp.369–379.

Ritter, W. E., & Johnson, M. E., 1911. The growth and differentiation of the chain of *Cyclosalpa affinis* Chamisso. Journal of Morphology, 22(2), 395–453.

Steinberg, D. K., Stamieszkin, K., Maas, A. E., Durkin, C. A., Passow, U., Estapa, M. L., … & Siegel, D. A., 2023. The Outsized Role of Salps in Carbon Export in the Subarctic Northeast Pacific Ocean. Global Biogeochemical Cycles, 37(1), e2022GB007523.

Sutherland, K. R., & Weihs, D., 2017. Hydrodynamic advantages of swimming by salp chains. Journal of The Royal Society Interface, 14(133), 20170298.

Sutherland, K. R., Gemmell, B. J., Colin, S. P., & Costello, J. H., 2019. Maneuvering performance in the colonial siphonophore, *Nanomia bijuga*. Biomimetics, 4(3), 62.

Tsagkogeorga, G., Turon, X., Hopcroft, R. R., Tilak, M. K., Feldstein, T., Shenkar, N., … & Delsuc, F. (2009). An updated 18S rRNA phylogeny of tunicates based on mixture and secondary structure models. BMC Evolutionary Biology, 9, 1–16.

Van Soest, R., 1974. Taxonomy of the subfamily Cyclosalpinae Yount, 1954 (Tunicata, Thaliacea), with descriptions of two new species. Beaufortia, 22(288), pp.17-55.

Yount J. L., 1954. The taxonomy of the Salpidae (Tunicata) of the Central Pacific Ocean. Pac. Sci. 8(3): 276–330

Yue, C. et al., 2015. Mechantronic system and experiments of a spherical underwater robot: SUR-II. J. Intell. Robot Syst. Doi:10.1007/s10846-015-0177-3.

