## Supplemental Table 1 for "A developmental ontology for the colonial architecture of salps"

| Species | Architecture | Adult colony specimens | Total adult colony specimens | Developing colony specimens | Total developing colony specimens |
|---|---|---|---|---|---|
| *Pegea sp.* | Transversal | **5** | **10** | **4** | **5** |
| *Pegea confoederata* | Transversal | **4 (Semenov, LJL videos)** | | | |
| *Pegea socia* | Transversal | **2 (R/V Sikuliaq 2022)** | | | |
| *Pegea bicaudata* | Transversal | **1 (Ryo Minemizu)** | | | |
| *Traustedtia multitentaculata* | Transversal | **1 (Dennis Rabeling)** | | **1 (Suzan Meldonian)** | |
| *Cyclosalpa affinis* | Whorl | **3** | **10** | **1** | **6** |
| *Cyclosalpa bakeri* | Whorl | **4** | | **2** | |
| *Cyclosalpa quadriluminis* | Whorl | **3** | | **3** | |
| *Cyclosalpa polae* | Cluster | **5** | **9** | **4** | **7** |
| *Cyclosalpa sewelli* | Cluster | **3** | | **3** | |
| *Cyclosalpa pinnata* | Cluster | **1 (Kona GoPro footage)** | | | |
| *Helicosalpa younti* | Helical | **1** | **2** | | **2** |
| *Helicosalpa virgula* | Helical | **1** | | **2 (D. Wrobel , Ringvold et al. 2020 Fig. 2EF)** | |
| *Helicosalpa komaii* | Helical | **1 (Ringvold et al. 2020 Fig. 4)** | | | |
| *Thalia cicar* | Oblique | **1** | **6** | **1 (KAUST formalin specimen)** | **2** |
| *Thalia longicauda* | Oblique | **2** | | | |
| *Thalia democratica* | Oblique | **1 (David Shale)** | | | |
| *Thetys vagina* | Oblique | **2 (NaturePL, MBA)** | | **1** | |
| *Iasis cylindrica* | Linear | **6** | **18** | **6** | **14** |
| *Ihlea punctata* | Linear | **1** | | **1** | |
| *Metcalfina hexagona* | Linear | **1** | | | |
| *Salpa aspera* | Linear | **1** | | | |
| *Salpa fusiformis* | Linear | **2** | | **4** | |
| *Salpa maxima* | Linear | **3** | | | |
| *Salpa thomspsoni* | Linear | **1 (LJL video)** | | **1 (Mike Stukel)** | |
| *Soestia zonaria* | Linear | **3** | | **2 (FSU photos)** | |
| *Brooksia rostrata* | Bipinnate | **3** | **9** | **2** | **6** |
| *Rittereilla amboinensis* | Bipinnate | **3** | | **3** | |
| *Ritteriella retracta* | Bipinnate | **3** | | **1** | |
